# Lymph Node Dissection and Radiation in the Rat Popliteal Region Leads to Progressive Lymphatic Pump Failure and Lymphedema

**DOI:** 10.64898/2026.03.11.711207

**Authors:** Shao-Yun Hsu, Young Jae Ryu, Zhanna Nepiyushchikh, Kiyoung Jeong, Jinseo Yu, J. Brandon Dixon

## Abstract

**Background:** Lymphedema, a chronic and incurable condition with limited therapeutic options, has limited options to quantitatively assess functional changes during its development; as a result, a deeper understanding of its pathophysiology remains hindered. To characterize lymphatic alterations and their association with disease pathology in a clinically relevant model in the rat, we developed a longitudinal iPhone-based volumetry method combined with non-invasive NIR analysis of lymphatic function.

**Material and methods:** Secondary lymphedema was induced by surgery and single-dose irradiation. iPhone volumetry provided longitudinal measurements of hindlimb volume, while NIR imaging quantified the pumping function of major lymphatic vessels in the popliteal area.

**Results:** Among 30 rats, lymphedema developed in 80%, defined as interlimb volume differences exceeding 5% and persisting through 14 days. In all rats with lymphedema, disease persisted until the end of the study at postoperative day 42 (P = 0.0015). NIR imaging revealed lymphatic dilation, dye extravasation, and lymphangiogenesis in affected limbs. Lymphatics in limbs with lymphedema exhibited increased contraction frequency, reduced amplitude, and diminished transport compared to baseline and contralateral controls (all P < 0.05). In contrast, rats that did no develop lymphedema showed no postoperative functional changes, although at baseline they displayed higher frequency and lower amplitude and transport compared with LE rats (all P < 0.001). Baseline transport values correlated negatively with swelling (r = –0.44, P = 0.002), as determined by ROC analysis, which yielded an AUC of 0.83, a sensitivity of 83.3%, and a specificity of 82.6%. Histopathology at day 42 confirmed significant dermal thickening and fat deposition in LE limbs (P < 0.001 and P = 0.002, respectively).

**Conclusion:** Longitudinal volumetry and NIR imaging applied to a clinically relevant animal model suggest a strong association between swelling and lymphatic function, which could provide deeper insight into lymphedema pathophysiology and represent valuable tools for future research and therapeutic development.

## 1. Introduction

Lymphedema arises from either congenital lymphatic underdevelopment or acquired damage to the lymphatic system, leading to excessive interstitial fluid and lipid accumulation within the affected limb.^1^ This pathological fluid retention disrupts immune homeostasis, increases susceptibility to recurrent infections, and contributes to progressive tissue fibrosis and limb deformity.^1–3^ Beyond physical impairment, lymphedema significantly reduces patients’ quality of life, mobility, and limb aesthetics. In severe cases, extensive vascular remodeling within the lymphedematous limb, in conjunction with immune dysfunction-induced hypercoagulability, may result in life-threatening complications such as stroke, pulmonary embolism, myocardial infarction, and, in extreme cases, mortality. ^1–7^ Despite ongoing research efforts, no definitive curative treatment for lymphedema currently exists, and disease management primarily relies on lifelong compression therapy to mitigate fluid and fat accumulation and slow disease progression.^8–10^ The most prevalent etiology of lymphedema is cancer treatment, particularly lymphadenectomy and radiation therapy, with an incidence rate of 30–35% among oncology patients. ^11–15^ Till 2022, lymphedema affects more than 35 million individuals in the United States and approximately 150 million worldwide, posing a substantial global healthcare challenge.^16^ The condition imposes significant economic and human resource burdens on healthcare systems, underscoring the necessity for prioritizing lymphedema management at both national and international levels. ^1,16^

The lymphatic system plays a critical role in maintaining interstitial fluid homeostasis, immune regulation, and lipid absorption.^17^ Lymphatic transport must overcome an adverse pressure gradient, as interstitial fluid pressure is generally lower than that of the venous system, to which lymph ultimately drains.^17–20^ Additionally, lymphatic fluid must be propelled against gravitational forces, particularly in the lower extremities. The primary mechanism facilitating lymphatic drainage is the intrinsic contractility of collecting lymphatic vessels (lymphangion), which contain specialized lymphatic muscle cells (LMCs) and unidirectional valves to ensure efficient fluid propulsion.^18,21^ LMCs are molecularly and functionally distinct from vascular smooth muscle cells, exhibiting characteristics common to both vascular and cardiac muscle.^22^ Dixon et al. demonstrated in a murine tail lymphedema model that lymphatic vessels progressively lose their contractility, ultimately failing to generate adequate pumping pressure.^23^ The severity of lymphedema progression correlates directly with the sustained loss of contractile function in the affected lymphatic vessels. ^23,24^ Understanding the functional changes of lymphatic vessels following injury is crucial for investigating the pathophysiology of lymphedema and developing advanced therapeutic strategies.^10,25,26^ However, prior studies examining lymphatic functional alterations have predominantly utilized tail models. In most models, the entire lymphatic drainage system in the tail is blocked, leaving no path for lymph outflow, a scenario that rarely ever occurs clinically. A deviation of this model in which one lymphatic outflow pathway in the tail is left intact overcomes this limitation. Yet, unlike in clinical lymphedema, the swelling will resolve on its own given time; therefore, functional analyses derived from the rodent tail model may not accurately recapitulate the dynamic lymphatic alterations that occur during the development of lymphedema in patients ^23,24,27,28^. Future research must refine experimental models in which lymphedema persists, even when lymphatic vessels are present in the limb, thereby enhancing clinical translatability and optimizing therapeutic development. ^23,24,27^ In 2014, Yang et al. developed the first rat hindlimb model of secondary lymphedema induced by a combination of surgical lymphatic disruption and radiation. ^29^ They systematically tested varying degrees and extents of lymphatic injury, along with different radiation doses, to optimize the balance between lymphedema incidence and animal mortality and complication rates. ^29^ Due to its high anatomical and etiological similarity to human lymphedema—and, most importantly, its ability to produce persistent edema that does not resolve spontaneously—this model has been widely adopted and refined over the past decade.^30–35^ Nonetheless, previous studies using this model have primarily focused on evaluating therapeutic strategies, including pharmacological and surgical interventions, and their effects on edema resolution and histological or immunological changes. ^30–35^ To date, no investigation has examined changes in lymphatic pumping function during the progression of lymphedema. Furthermore, because most studies have relied on CT or MRI to measure hindlimb volume, volume monitoring has typically been limited to only two or three time points, lacking longitudinal and dynamic observation. ^30–35^

Furthermore, the effectiveness of lymphedema treatment is highly dependent on the timing of intervention.^10,25,26^ Early diagnosis and treatment have been associated with significantly superior clinical outcomes. ^10,12,25^ Nevertheless, current diagnostic practices often fail to detect lymphedema until lymphatic vessel pumping dysfunction has already resulted in overt edema, thereby missing the critical window for early intervention.^10,25^ Clinical studies have shown that preventive lymphovenous anastomosis (LVA) can reduce the incidence of developing lymphedema in breast cancer patients. ^36,37^ Nonetheless, the lack of validated methodologies to identify individuals with intrinsically vulnerable lymphatic vessels limits the precision of patient selection, thereby attenuating the preventive efficacy of such procedures. ^12,36,37^ In addition, considering that prophylactic surgical interventions may pose a risk of lymphatic injury, their application in patients with a low risk of lymphedema may conflict with the principle of no-harm. ^13,15,26,36–38^ The absence of reliable tools to distinguish high-risk from low-risk individuals prior to disease onset remains a significant obstacle to the broader implementation of preventive strategies.^26,36,37^

This study aims to address these critical gaps by utilizing a non-invasive, quantitative method to assess lymphatic functionality in both healthy and diseased states within a clinically relevant secondary lymphedema rat model. Through this research, we aim to elucidate the changes in lymphatic pumping function during the progression of secondary lymphedema, thereby enhancing the understanding of lymphedema pathophysiology and identifying early disease markers that may facilitate timely intervention and improved therapeutic strategies.

## 2. Material and Methods

### 2.1. Animal Model

The animal study was approved by our Institutional Animal Care and Use Committees (IACUCs) (IACUC protocol number: XXX A100297). Sixteen-week-old female Lewis rats, an inbred strain recognized for its suitability in transplantation studies, were utilized in this study. The rat model was designed to simulate the clinical presentation of secondary lymphedema. This was achieved through the surgical excision of major inguinal lymphatic vessels, the entire inguinal fat pad containing all inguinal lymph nodes, and the popliteal fat pad containing all popliteal lymph nodes via an oblique inguinal incision on the right hindlimb. On postoperative day four, the right hind limb was subjected to localized irradiation under general anesthesia. The right hindlimb of the rats was irradiated using an RS-2000 small animal X-ray irradiator (Rad Source Technologies©). A total dose of 22.4 Gy was administered at a rate of 1 Gy per minute, with an operating voltage of 225 kV and a current of 12 mA. A custom-designed lead shield was employed to ensure targeted exposure, protecting all body regions except the irradiated hindlimb. The contralateral limb was left unmanipulated and served as an internal control.

Lymphedema was diagnosed when the volume of the affected limb exceeded that of the contralateral limb by more than 5%, as assessed two weeks post-irradiation. Limb volume was quantified using an iPhone-based volumetric analysis protocol developed in-house, employing the formula: (Volume of the affected limb − Volume of the healthy limb) / Volume of the healthy limb × 100%.) The current study is conducted and reported in line with the ARRIVE guideline (Standards for the Reporting of Experimental Animal Studies).

### 2.2. Grouping and Timeline

Prior to the surgical procedure, the volumes of both hindlimbs of the rats were measured using iPhone Volumetry. Only those rats exhibiting a volume difference of less than 5% between the contralateral limbs were included in the study. Of the 30 rats assessed, 29 (96.7%) met this criterion. For these selected animals, near-infrared (NIR) imaging of popliteal lymphatic vessels in both limbs was performed. Following the imaging, the surgical procedure was carried out. On postoperative day 4, the right hindlimb was irradiated. Following surgery and irradiation, the volumes of the hindlimbs were assessed twice weekly, and vessel functionality was evaluated biweekly using NIR imaging. Fourteen days post-irradiation, rats in which the volume of the right hindlimb exceeded that of the left hindlimb by more than 5% were categorized into the lymphedema group. Rats that did not meet this criterion were classified into the non-lymphedema group.

### 2.3. iPhone volumetry

The iPhone 13 Pro (Apple Inc.) was used to capture images of the rats following the sample code “Taking Pictures for 3D Object Capture (Apple Inc.)”^39^ released by Apple. According to Apple’s guidelines, each capture session consisted of at least 30 individual photos to cover both hindlimbs of the rat comprehensively. The sample code recorded the image of each capture, the distance between the camera and the rat, and the camera’s tilt angle. These data were stored in a folder named according to the timestamp of the first capture. Subsequently, a MacBook Pro (Apple Inc.) with an Apple M1 Pro chip, 16GB memory, 6400 MHz LPDDR5, and a 1TB SSD was used to run the Apple-released sample code “Creating a Photogrammetry Command-Line App (Apple Inc.)”^40^ to convert the recorded data into a 3D point cloud model. The 3D point cloud model was then imported into Blender 3.4.1 (Blender Foundation) for processing. The model was refined using a re-mesh with 8 octree depth, 0.90 scale, and a 1.00 threshold. The volume of the rat’s hindlimbs was calculated separately on each side by using the inguinal crease to segment the limb.

### 2.4. NIR Image Functional Analysis

Rats were placed on a heating pad to maintain their body temperature. A total of 20 µL of PEG-IRDye (methods previously described ^41^) was injected intradermally approximately 1 inch upstream from the paw tip. Three minutes after injection, dynamic lymphatic transport was then captured at a rate of 4 frames per second with a 150 ms exposure time over a period of 13 minutes on the medial side of the popliteal area using a near-infrared lymphatic imaging system.^42^ The system involves an Evolve MVX-ZB10 microscope (Olympus), a Lambda LS xenon arc lamp (Shutter Instruments, a 769-nm bandpass excitation filter, an 832-nm emission bandpass filter, and an Evolve Delta 512 ECMCCD camera (Photometrics).

Evaluation of lymphatic function from NIR videos was performed by first importing the entire sequence into ImageJ, version 1.53K (NIH, U.S. Government), and calculating the standard deviation of intensity variability for each pixel. From these data, five regions of interest (ROIs) along the major lymphatic vessel with the most prominent intensity changes were selected for analysis. Using these ROIs, three distinct functional metrics were subsequently calculated using established methodologies in MATLAB R2023a (MathWorks) methods, as previously described in detail. ^43^ In brief, packet frequency was determined by identifying all fluorescence intensity peaks within the ROI time series and measuring the time intervals between consecutive peaks. The reciprocal of each interval was calculated to represent the instantaneous contraction rate, and these values were averaged to obtain the overall frequency of lymphatic contractions during the imaging period. (Supplement 1)

Packet amplitude was assessed by measuring the difference in fluorescence intensity between each peak and its preceding trough within the same cycle. This difference reflects the magnitude of fluorescence fluctuation associated with vessel contraction, and the mean amplitude across all cycles was used as the representative value for each ROI. (Supplement 1)

Because in NIR imaging the intensity peak corresponds to the vessel being fully dilated with the maximum amount of dye in the lumen, while the lowest intensity occurs when the vessel is maximally contracted and has transported dye to the next segment, packet transport was evaluated by integrating the fluorescence signal along the curve for each packet. This approach captures the cumulative change in fluorescence intensity throughout the contraction cycle, providing a measure of how much tracer is effectively moved during each pumping event. The resulting value was normalized by the duration of the packet to account for differences in cycle length. (Supplement1)

### 2.5. Histology

After euthanasia, 5 × 5 mm skin samples were harvested bilaterally from the ventral thigh of five rats, including the skin and all connected tissues above the bone. Samples were fixed in formalin, embedded in optimal cutting temperature (OCT) compound, and stored at −80 °C. Frozen tissues were subsequently cryosectioned in 20 um thickness and mounted on glass slides for staining and imaging. Hematoxylin and eosin (H&E) staining was performed using a Leica Autostainer XL (Leica Biosystems). For each leg, four sections were stained. Images were acquired at 10 times magnification using a Zeiss Axio Observer Z1 microscope (Zeiss). Dermal and adipose thickness were blindly quantified manually with ImageJ software (version 1.53K, NIH, U.S. Government) by tracing the visible dermal and fat regions in H&E-stained sections. Thickness was calculated by dividing the total traced area of dermis or fat by the corresponding sample width.

### 2.6. Statistics

Descriptive statistics for continuous variables included the mean, standard deviation, and range, while categorical variables were summarized using frequencies and percentages. Nested t-tests, two-sample t-tests, or Mann-Whitney U tests were applied as appropriate to compare two independent means. Statistical significance were defined at a p-value of 0.05. Receiver operating characteristic (ROC) analysis was employed to assess the accuracy of functional features in predicting the occurrence of lymphedema. All statistical analyses were performed using Prism 10 (GraphPad LLC). Consistency of iPhone-based volumetric measurements was assessed in R by calculating the mean coefficient of variation (CV) across repeated measurements, providing a quantitative measure of reproducibility.

## 3. Results

### 3.1. Longitudinal Volume Changes

By day 14, lymphedema developed in 80% (24/30) of rats that underwent lymph node dissection followed by a single dose of irradiation. From day 0 to day 14 post-irradiation, the volume difference between the lymphedema leg and the normal leg was significantly greater than the preoperative baseline. (Preoperative baseline: mean ± SD = -0.065 ± 3.16%; day 0: 15.80 ± 16.54%, P < 0.001; day 3: 26.84 ± 35.24%, P < 0.0001; day 7: 27.36 ± 23.12%, P < 0.0001; day 10: 22.92 ± 19.67%, P < 0.0001; day 14: 23.76 ± 11.42%, P < 0.0001; Figure 1D). Combined lymph node dissection and irradiation resulted in sustained lymphedema, with swelling persisting throughout the observation period. At 42 days postoperatively, rats with lymphedema continued to exhibit significantly greater interlimb volume differences compared to baseline (preoperative: mean ± SD = -0.065 ± 3.16%; day 42: -0.94 ± 2.65%, P <0.01), with a persistent increase exceeding 5% (Figure 1E) iPhone volumetry measurements showed high reproducibility, with 80% of measurements having a CV below 0.032 and the 99th percentile remaining under 0.045, indicating minimal variability across repeated assessments. (Figure 2)

**Figure 1.**
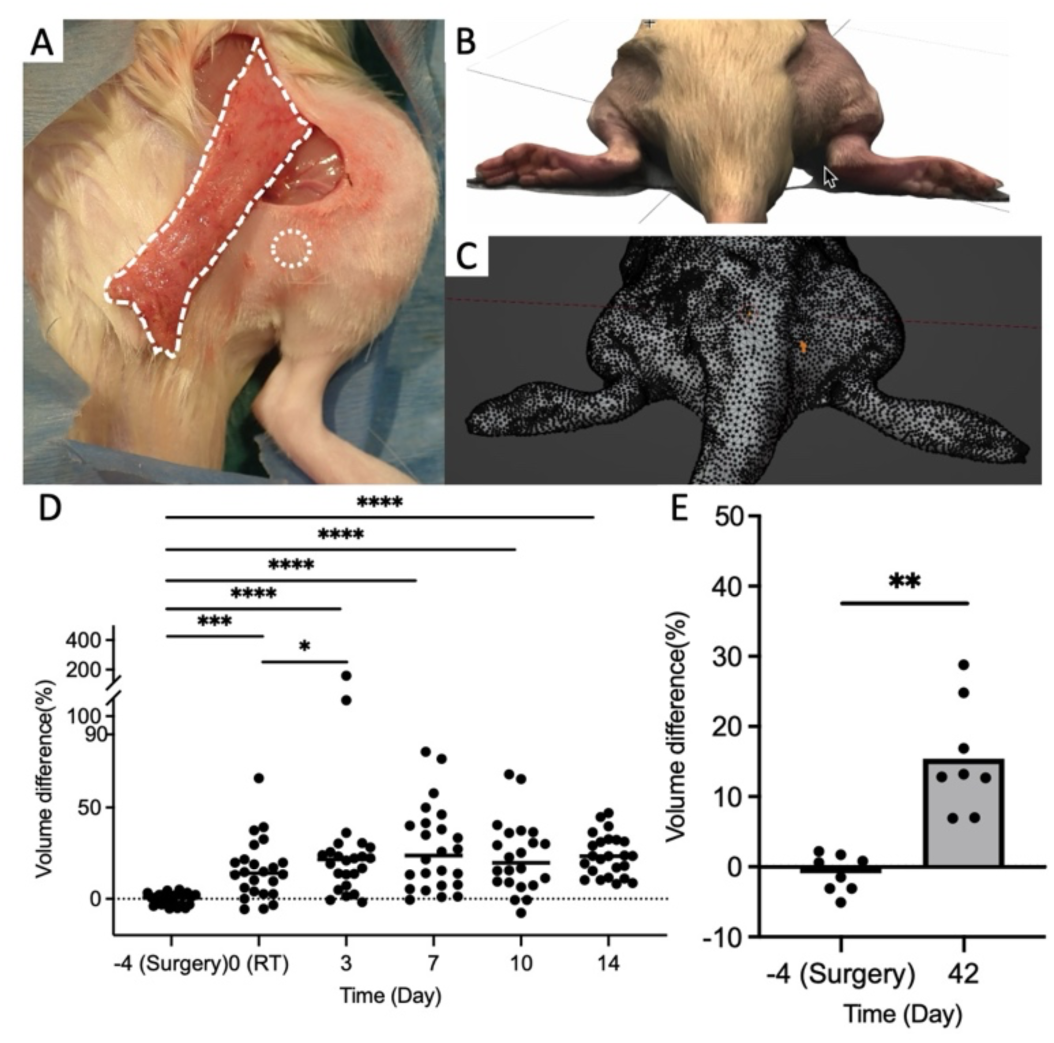
Lymphedema rat model. (A) Lymphedema is created by removing the popliteal and inguinal LNs with the afferent and efferent LVs supplying the LNs on one side of the rat. 3 days after surgery, the site is irradiated with 22.7 Gy, (B) Photographic image, and (C) 3D image reconstruction from point cloud. The swelling started 4 days after irradiation (D) and persisted for 42 days after the irradiation.

**Figure 2.**
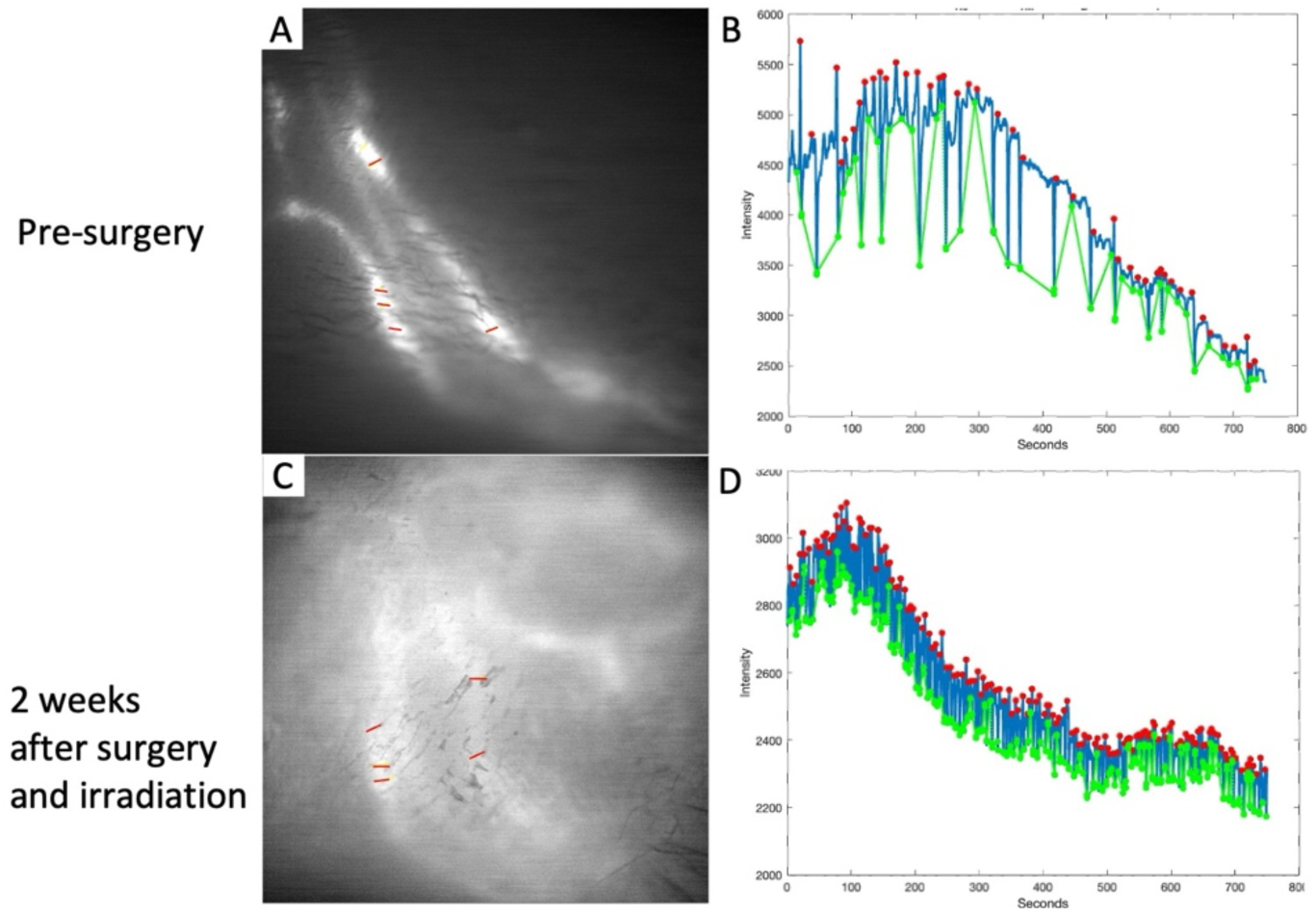
Distribution of Coefficient of Variation (CV) and summary statistics at selected percentiles. The histogram illustrates the distribution of CV values across samples, with most values concentrated in the lower range and a right-skewed tail extending toward higher CV values. The accompanying table summarizes the mean CV (%) at different percentile thresholds: 80% (3.23), 95% (4.06), 99% (4.42), and 100% (4.54).

### 3.2. Lymphatic Structural and Functional Alterations

In NIR imaging, the LE limb exhibited marked lymphatic dilation, dye extravasation, and evidence of lymphangiogenesis (Figure 3C, Supplementary Video 2) compared to the pre-surgical baseline. (Figure 3A, Supplementary Video 1.) A representative tracing of temporal intensity changes in the most dynamic ROIs demonstrated that the LE limb displayed more frequent but smaller amplitude fluctuations in signal intensity (Figure 3D). Averaging five ROIs per animal revealed that, at two weeks post-irradiation, the popliteal lymphatic vessels of the lymphedema legs exhibited significantly increased contraction frequency (baseline: 6.41 ± 2.35; 2 weeks: 11.67 ± 4.51), reduced contraction amplitude (baseline: 0.22 ± 0.12; 2 weeks: 0.11 ± 0.08), and decreased lymphatic transport (baseline: 36.37 ± 11.46; 2 weeks: 23.17 ± 14.13), compared with baseline (Figures 4A–C; P < 0.0001 for all). A similar pattern was observed when comparing LE limbs with contralateral normal limbs at the two-week time point; LE limbs demonstrated significantly higher pumping frequency (LE: 11.67 ± 4.51 ; normal: 8.881 ± 8.423, P = 0.01), lower pumping amplitude (LE: 0.11 ± 0.08; normal: 0.17 ± 0.14, P <0.01), and diminished pumping transport (LE: 23.17 ± 14.13; normal: 31.06 ± 24.36, P = 0.02) (Figures 4D–E).

**Figure 3.**
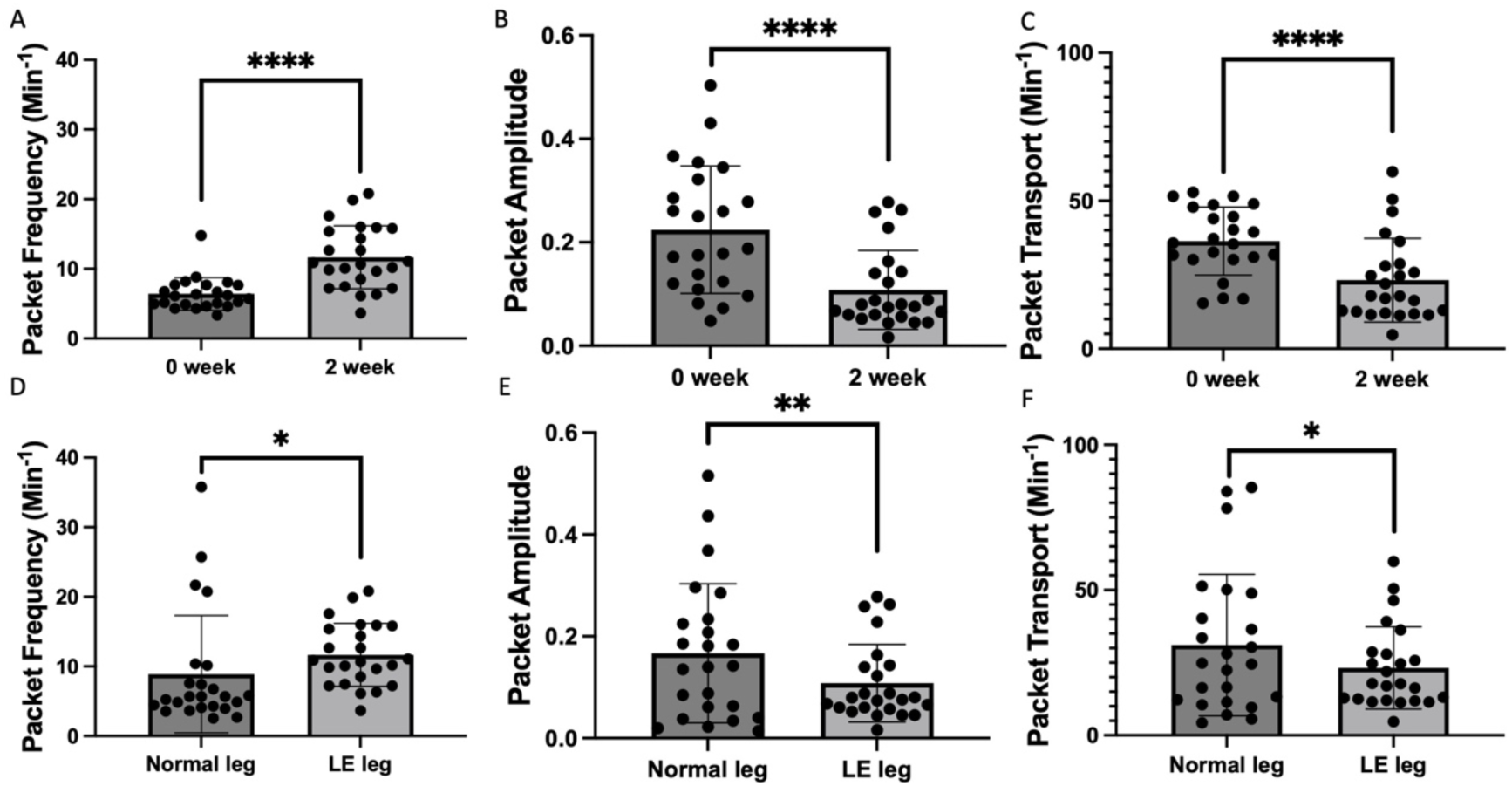
Lymphatic vessels in the lymphedema limb show structural and functional disruption two weeks after the surgery and irradiation, characterized by (C) dilation, leakage, and regenerative sprouting suggestive of compensatory remodeling compared to a) pre-surgery NIR image, and (B), (D) quantification of intensity fluctuations from lymphatic contraction from videos of 1 cm proximal to the ankle, as in (A), AC), retrospectively. The red strips mark the selected ROIs.

**Figure 4.**
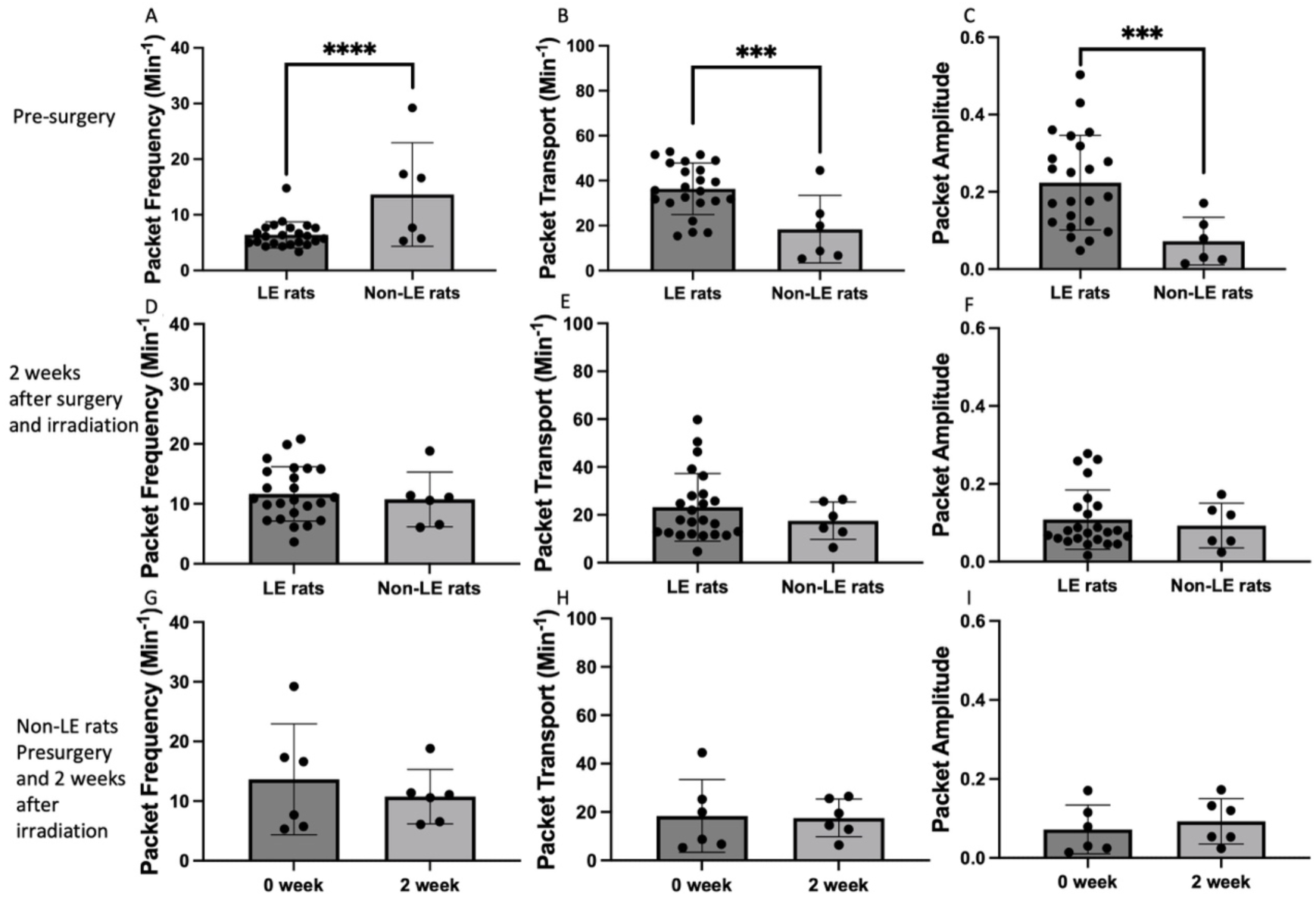
Lymphatic function is reduced after injury in the affected limbs. This is reflected in (A) an increased packet frequency, (B) decreased packet amplitude, and (C) decreased packet transport in lymphedema rats 2 weeks after irradiation within the edematous limb. The same significant differences in (D) packet frequency, (E) packet amplitude, and (F) packet transport are found when comparing the edematous limb with the contralateral (normal leg).

### 3.3. Lymphatic Function in Non-LE Rats

Using the same NIR imaging and quantitative analysis, we compared lymphatic functional changes between LE rats and rats that underwent identical lymph node dissection and irradiation but did not develop lymphedema (non-LE rats). At baseline (day−4, pre-surgery and pre-irradiation), non-LE rats exhibited significantly higher pumping frequency (non-LE: 13.65 ± 9.30; LE: 6.41 ± 2.35, P < 0.0001), lower pumping amplitude (non-LE: 0.07 ± 0.06; LE: 0.22 ± 0.12, P < 0.001), and lower pumping transport (non-LE: 18.41 ± 15.07; LE: 36.37 ± 11.46, P < 0.001) compared with LE rats (Figures 5A–C). At two weeks post-irradiation, no significant differences were observed between non-LE and LE rats in pumping frequency (non-LE: 10.75 ± 4.58; LE: 11.67 ± 4.51, P = 0.39), amplitude (non-LE: 0.09 ± 0.06; LE: 0.11 ± 0.08, P = 0.38), or transport (non-LE: 17.54 ± 7.79; LE: 23.17 ± 14.13, P = 0.10) (Figures 5D–E). Furthermore, in non-LE rats, these functional measurements at two weeks postoperatively did not differ significantly from their baseline values (baseline: 13.65 ± 9.30; two weeks: 10.75 ± 4.58 for pumping frequency, P = 0.18; baseline: 0.07 ± 0.06; two weeks: 0.09 ± 0.06 for amplitude, P = 0.26; baseline: 18.41 ± 15.07; two weeks: 17.54 ± 7.79 for transport, P = 0.80) (Figures 5 G-I).

**Figure 5.**
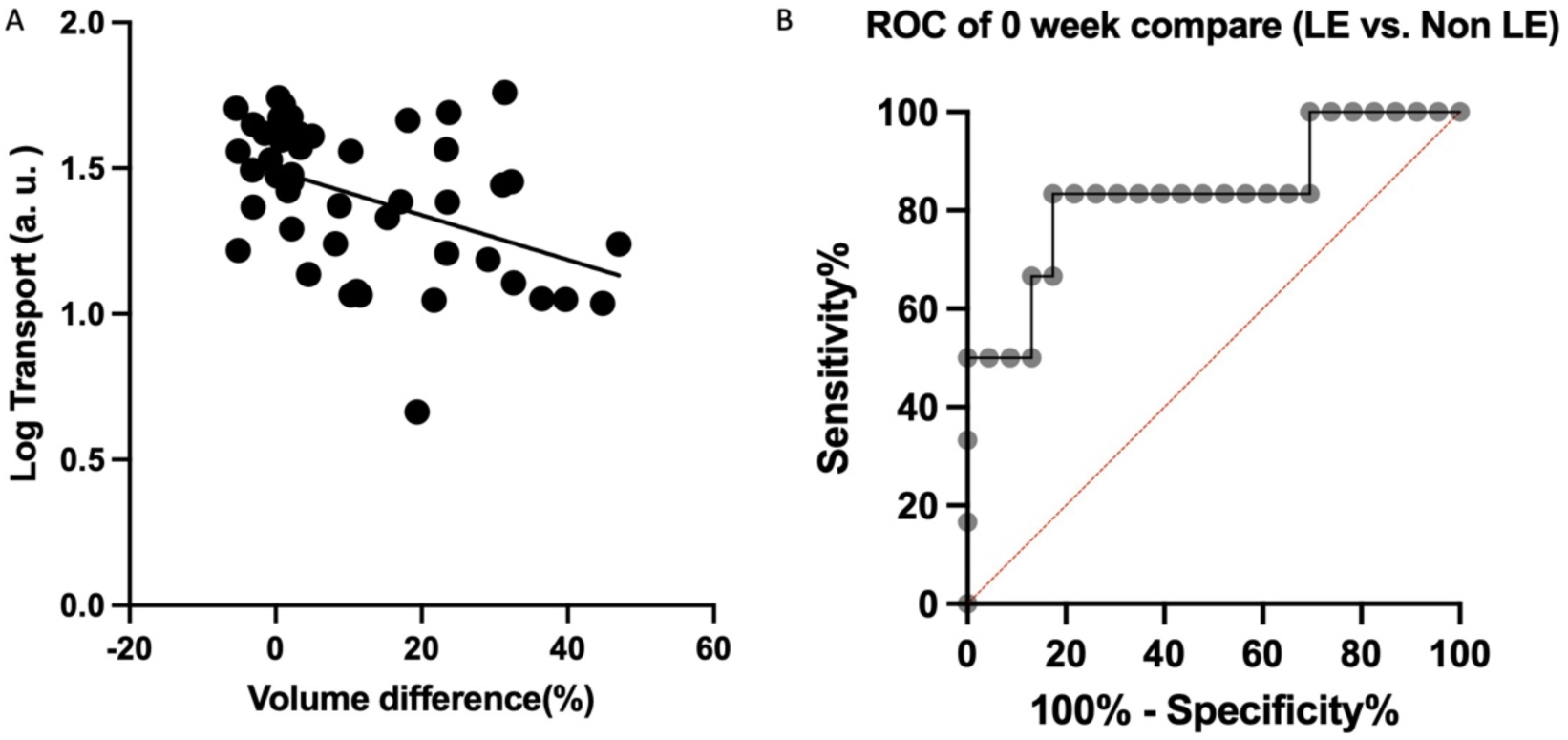
Non-LE rats exhibit distinct baseline lymphatic function that may protect against the development of lymphedema. This is reflected in (A) higher packet frequency, (B) lower transport, and (C) reduced amplitude before surgery compared with rats that later developed LE. Two weeks after surgery and irradiation, no significant differences in these functional parameters were observed between LE and non-LE rats (D), (E), and (F), suggesting that non-LE rats may possess additional peripheral drainage pathways and thus rely less on the single popliteal-to-inguinal route. Before and after surgical intervention and irradiation, significant alterations were observed in packet frequency, packet amplitude, and packet transport in LE rats (A), (B), and (C). In contrast, non-LE rats demonstrated no significant differences across the same parameters (G), (H), and (I), indicating that the lymphatics of non-LE rats are less susceptible to the detrimental effects of surgical injury and irradiation compared to those of LE rats.

### 3.4. Predictive Value of Baseline Lymphatic Transport

Analysis of lymphatic functional changes (pumping transport) in relation to lymphedema severity revealed a significant negative correlation. Log-transformed transport values were inversely correlated with interlimb volume differences (Figure 6A; P = 0.002, R = −0.4446). Furthermore, ROC analysis of baseline transport values in LE and non-LE rats demonstrated high discriminative power, with an area under the curve (AUC) of 0.833. The optimal cutoff value was 27.68, yielding a sensitivity of 83.33% and a specificity of 82.61% (Figure 6B)

**Figure 6.**
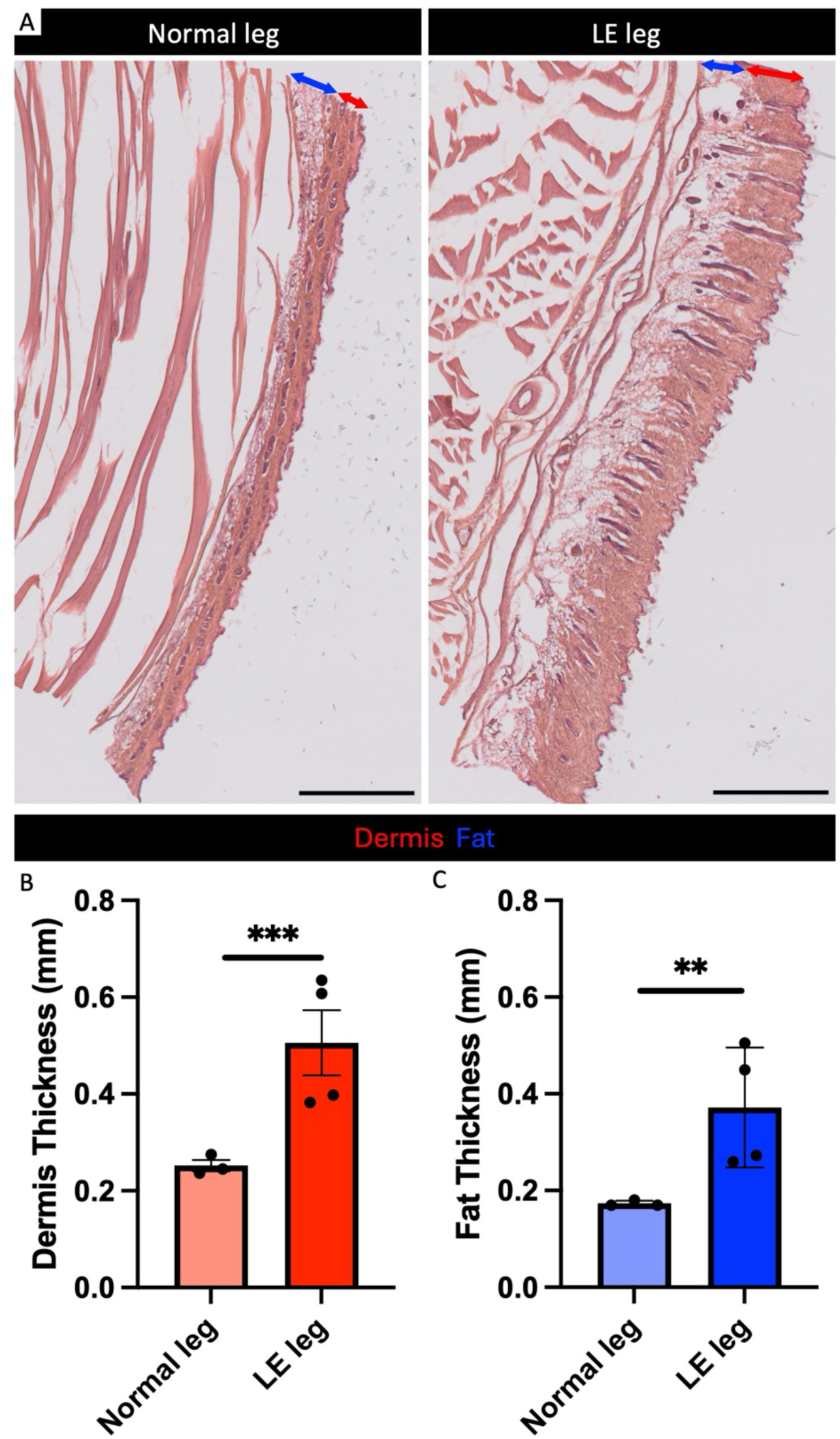
Lymphatic transport as measured by NIR, negatively correlates with swelling, and is predictive of postoperative lymphedema development. (A) A significant negative correlation was observed between log-transformed transport values measured by NIR and volume differences (P = 0.002, R = –0.4446). (B) Preoperative transport values demonstrated good predictive performance for postoperative lymphedema, with ROC analysis showing an AUC of 0.833, sensitivity of 83.33%, and specificity of 82.61%.

### 3.5. Histopathological Features of Lymphedema

Histological analysis revealed that, 42 days after irradiation, the LE limb exhibited thickening of both the dermis and subcutaneous fat layers. Quantitative image analysis confirmed that the LE limb had significantly greater dermal thickness (LE: 0.51 ± 0.13 mm; normal: 0.25 ± 0.02 mm, P < 0.001) and subcutaneous fat deposition (LE: 0.37 ± 0.12 mm ; normal: 0.17 ± 0.01, P <0.01) compared with the contralateral normal limb (Figures 7A–C).

**Figure 7.**
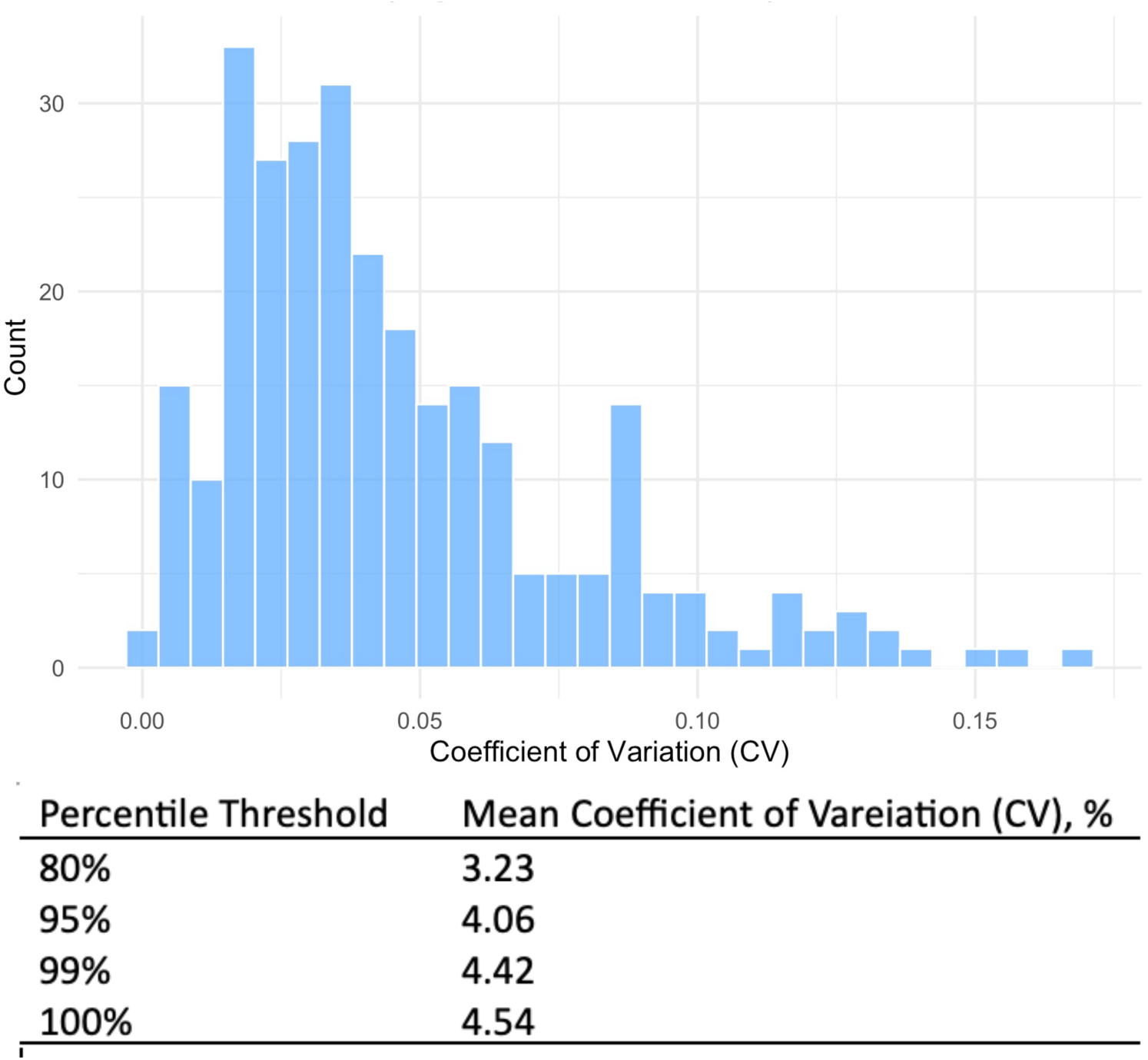
Dermal and subcutaneous adipose layers are significantly thickened in the LE leg compared with the normal leg. (A) Representative histological images show marked expansion of the dermis (red) and subcutaneous fat (blue) in the LE leg relative to the normal leg. (B) Quantitative analysis confirms a significant increase in dermal thickness, and (C) a significant increase in subcutaneous fat thickness in the LE leg compared with the normal leg. Scale bars: 1 mm.

## 4. Discussion

Lymphedema remains an incurable condition, and animal models are essential for advancing therapeutic strategies.^1–3,8^ The mouse tail model is commonly used for drug screening due to its low cost; however, its anatomical limitations render it unsuitable for studying or advancing lymphatic reconstructive microsurgery, which is regarded as the most promising treatment approach.^27,28,44,45^ To overcome these limitations, the rat hindlimb model, which combines lymphadenectomy and irradiation, was developed to better replicate human anatomy, result in permanent irreversible lymphedema when no therapy is applied, and enable microsurgical interventions, becoming the most clinically relevant model over the past decade.^29,30,34,46,47^ Despite its advantages, assessment methods for this model have been limited primarily to volumetric measurements using computed tomography or magnetic resonance imaging, leaving the dynamic progression of swelling largely unexplored. ^29,30,34,46,47^ Furthermore, functional changes in lymphatics during lymphedema progression have not been investigated. ^29,30,34,46–48^ In this study, we introduced an integrated approach utilizing iPhone volumetry, NIR imaging for lymphatic function analysis, and quantitative histology analysis, enabling high-resolution, longitudinal assessment of swelling and quantitative evaluation of lymphatic pumping and tissue remodeling.

The lymphatic system shares several anatomical and functional features with the cardiovascular system;^17,18,49^ however, its much smaller scale precludes direct measurement of parameters such as flow rate, intramural pressure, and pumping pressure, which are routinely assessed in the cardiovascular system. ^17,18,50^ Currently, no clinical method exists for quantifying human lymphatic pumping function, and most previous in vitro studies have focused on physiological conditions or comprehensible injury rather than lymphedema states.^42,51–55^ Although Dixon et al. assessed impaired lymphatic function in mouse tail and sheep hindlimb models, these injuries were mild and fully compensated without persistent swelling, failing to replicate the irreversible dysfunction observed in lymphedema.^23,56,57^ In the present study, we longitudinally evaluated lymphatic pumping in an animal model that not only reflects clinical relevance but also fulfills diagnostic criteria for lymphedema. Following disease onset, lymphatic vessels exhibited more frequent but weaker contractions, a pattern analogous to congestive heart failure, where increased contraction frequency initially compensates for elevated preload and afterload but eventually leads to muscle fatigue and pump failure. This mechanistic similarity suggests that therapeutic strategies used in heart failure, such as rate control, may be applicable to preserving lymphatic function and preventing disease progression.^49,58,59^ Supporting this concept, Sestito et al. demonstrated that lymphatic-specific lipid nanoparticles delivering calcium channel agonists induced slower yet stronger lymphatic contractions, paralleling the benefits of heart rate control.^42^ Our findings provide further evidence for exploring rate-modulation approaches as a potential intervention for lymphedema.

Through NIR imaging and longitudinal volumetric assessment, we were able to investigate the relationship between lymphatic functional changes and the development of lymphedema. Previous studies examining pumping frequency have reported inconsistent findings: some observed an increase after lymphatic injury, while others reported a decrease.^23,24,28,56,57,60,61^ These discrepancies likely reflect methodological differences, as most studies used lymphatic injury models rather than clinically relevant lymphedema models. Consequently, pumping frequency alone does not reliably correlate with the degree of limb swelling. In contrast, packet transport—a quantitative measure representing the relative volume of lymphatic fluid transported by a selected lymphatic vessel per unit time—provides a more robust functional metric, as evidenced by the ability of baseline measurements of this parameter to reasonably predict the risk of lymphedema development (Fig. 5). This finding suggests that non-invasive functional imaging holds promise for early identification of patients at high risk for severe lymphedema by detecting early declines in lymphatic pumping capacity. In addition, prior cross-species studies have consistently demonstrated that transport decreases following lymphatic injury, aligning with our findings. ^23,24,28,56,57,60,61^

Beyond functional metrics, the observed association between lymphatic function and swelling severity offers new insights into the pathophysiology of lymphedema. Despite identical surgical and radiation protocols, 30–40% of patients develop lymphedema, and no current treatment is as effective as methods aimed to prevent its onset.^9,11,13,15^ Prophylactic LVA has been shown to reduce incidence by approximately 60%. Nevertheless, performing prophylactic LVA universally is neither practical nor ethical because identifying suitable lymphatic–venous pairs intraoperatively remains challenging, and performing the procedure on all patients would subject 60–70% of low-risk individuals to unnecessary surgical risk.^36–38^ In our study, rats subjected to the same surgery and irradiation exhibited an 80% lymphedema incidence. Interestingly, non-LE mice (those that did not develop lymphedema) exhibited inherently different lymphatic function compared to LE rat, and their pumping parameters remained stable before and after intervention (Figure 4g–i). Surprisingly, prior to intervention, non-LE rat already demonstrated lower baseline functional values compared to LE rat; however, these values did not significantly differ from those observed in LE rats two weeks after irradiation (Figure 4a–f). These observations are similar to those in certain patients, where an elevated measurement of lymphatic activity prior to surgery, was suggestive of higher lymphedema risk after surgery. ^62,63^

This study has several limitations. Although this rodent model is the most clinically relevant available, its small body size creates major differences in the impact of gravity on lymphatic function and remodeling compared with humans, which is likely why in the rat a relatively severe injury to the lymphatic system is necessary to induce lymphedema. Our NIR image analysis relied on selecting five ROIs along the popliteal major lymphatics with the most prominent intensity changes. Although we applied a standard deviation–based method to reduce subjectivity, operator-dependent bias could not be completely eliminated. Moreover, this approach evaluates only major lymphatic vessels, excluding smaller branches and intradermal lymphatics, which may explain why non-LE rats exhibited lower calculated drainage and function than LE rats, yet did not develop limb swelling, suggesting alternative drainage routes that were undetected by our method. Future machine learning–based analyses may overcome these shortcomings by enabling unbiased ROI selection and whole-image quantification. Nevertheless, the longitudinal iPhone-based volumetry and non-invasive NIR functional analysis techniques developed here provide more accessible and accurate methods for evaluating lymphatic pathophysiology and testing new treatment strategies. Furthermore, these tools, applied to a clinically relevant rodent model, our findings provide important insights into lymphedema pathophysiology, demonstrating how lymphatic function evolves through injury, remodeling, compensation, and eventual failure, and establishing correlations with limb volume increase. These results not only support the proposed similarity between lymphatic and cardiovascular pump failure but also provide a foundation for early, non-invasive diagnostic strategies and the development of novel treatments.

## 5. Conclusion

In summary, this study establishes a clinically relevant experimental platform that combines longitudinal iPhone-based volumetry with non-invasive NIR functional analysis to provide a practical and accurate assessment of lymphatic function. Using this approach, we delineated the dynamic progression of lymphatic remodeling from injury through compensation to failure, and demonstrated its close correlation with limb volume changes, the primary clinical manifestation of lymphedema. These findings not only reinforce the mechanistic parallels between lymphatic and cardiovascular pump failure but also highlight new opportunities for developing early diagnostic strategies and innovative therapeutic interventions for lymphedema.

## Ethical approval

The study was approved by the Institutional Animal Care and Use Committees

## Patient or volunteer consent

Not applicable

## Data statement

All data acquired and/or analyzed during this study are available from the corresponding author upon reasonable request.

## Appendix A. NIR Analysis Code

clear;

close all;

% -------------------------------------------------

% User-defined detection thresholds

%--------------------------------------------------

indicesThreshold = 0.001; % Threshold for defining minima boundaries (relative intensity tolerance)

minThreshold = 0.01; % Minimum allowed local prominence threshold for peak detection

sensitivityThreshold = 0.5; % Sensitivity for selecting optimal moving window size (fraction of tolerance)

% Consolidate thresholds into one vector for convenience

thresholds = [indicesThreshold; minThreshold; sensitivityThreshold];

% ------------------------------------

% Directory and output file setup

% -----------------------------------

myDir = uigetdir; % Prompt user to select input directory containing CSV files

myFiles = dir(fullfile(myDir,‘*.csv’)); % Identify all .csv files in the directory

% Create an output directory labeled with date/time stamp

FileName = sprintf(‘FunctionalAnalyses_UpsideIntegration_BatchOutput_%s’, datestr(now));

FileName = regexprep(FileName, ‘-’, ‘_’); % Sanitize filename (replace illegal characters)

FileName = regexprep(FileName, ‘ +’, ‘_’);

FileName = regexprep(FileName, ‘:’, ‘_’);

mkdir(sprintf(FileName))

FullFilePathName = fullfile(pwd, FileName);

% -------------------------

% Analysis settings

% -----------------------

PlotFigs = ‘y’; % Flag to plot and save figures for each file

Results = []; % Container for aggregated results

%-----------------------------------------

% Batch process each .csv file

% ----------------------------------------

e = 0;

k = 1;

while k <= length(myFiles)

baseFile1Name = myFiles(k).name;

Name = string(baseFile1Name(1:end-4)); % Strip extension for labeling

try

% Run packet frequency analysis with upside integration method

tempResult = PacketFreqUpside_AUTO([myDir ‘/’ baseFile1Name],

thresholds, Name, PlotFigs);

e = 0;

catch

% If analysis fails, fill with missing values

e = e + 1;

end

% Append results (successful or missing)

if e==0

Results = [Results; [Name tempResult]];

else

Results = [Results; [Name [missing missing missing missing missing

missing missing missing]]];

end

% Save figure output if enabled

if PlotFigs == ‘y’

figname = strcat(Name, ‘.fig’);

savefig(gcf, strcat(FullFilePathName,‘/’, figname));

end

k = k+1;

end

% Save compiled results as MATLAB .mat file

save([FullFilePathName ‘/Results’],‘Results’)

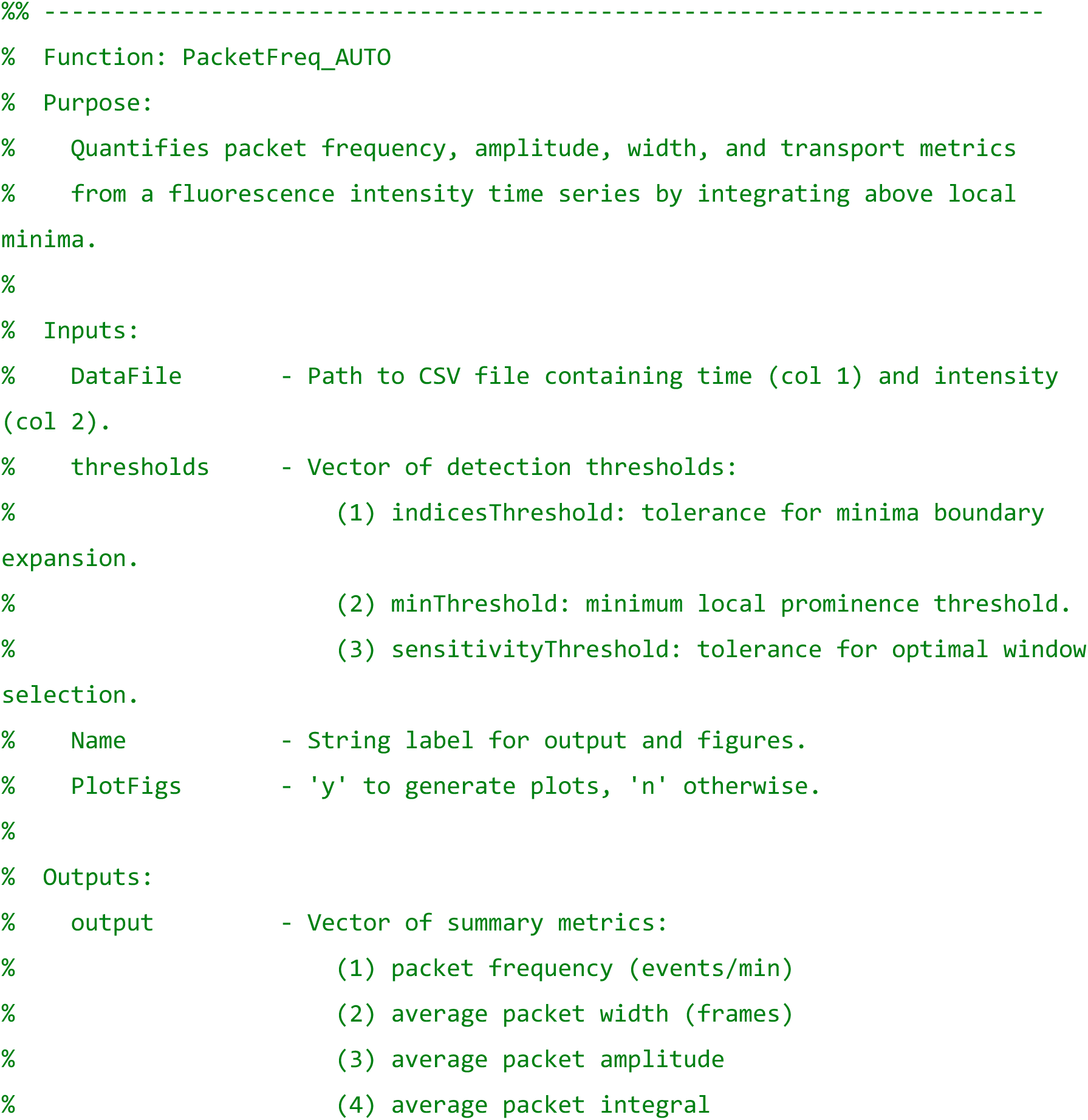
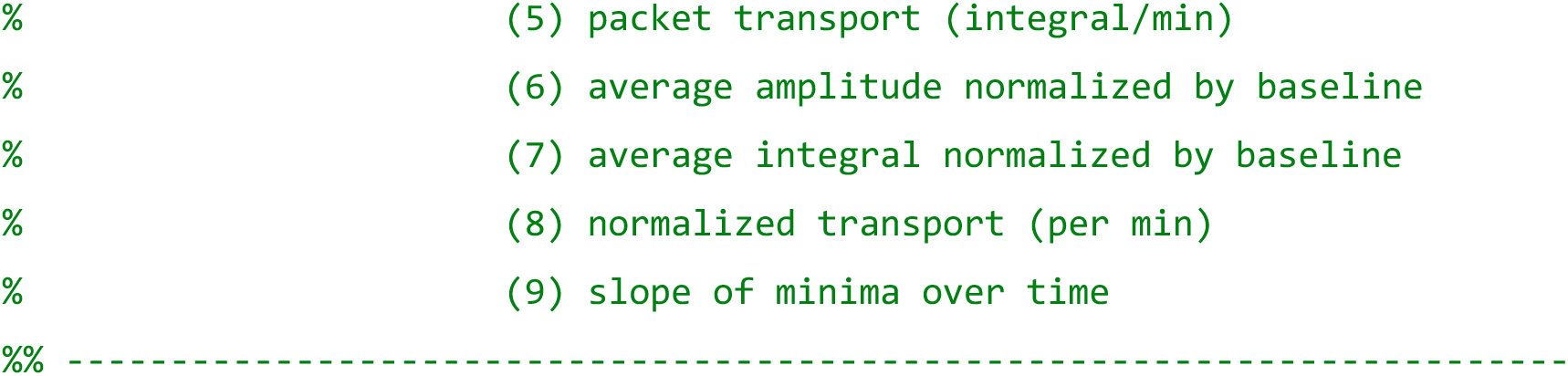

function output = PacketFreq_AUTO(DataFile, thresholds, Name, PlotFigs)

indicesThreshold = thresholds(1);

minThreshold = thresholds(2);

sensitivityThreshold = 1 - thresholds(3);

% Load CSV (time, intensity)

x = csvread(DataFile);

t = x(:, 1);

x = x(:, 2);

% Smooth trace (3-point moving average)

y2 = smooth(x,3);

% Determine optimal detection window size adaptively

numSoln = [];

minWindow = 1;

maxWindow = length(y2);

for i = minWindow:maxWindow

[x,y] = peakdetYJ(y2, i, minThreshold);

numSoln = [numSoln; length(x)];

end

windowSize = (minWindow:1:maxWindow-1)’;

DiffEQN = movmean(diff(numSoln),10);

OptWindowSize = find(abs(DiffEQN) <= sensitivityThreshold, 1);

% Peak detection with optimal window

[maxtab2, min2] = peakdetYJ(y2, OptWindowSize+1, minThreshold);

% Align maxima and minima

if maxtab2(1,1) < min2(1,1), maxtab2 = maxtab2(2:end,:); end

if maxtab2(end,1) > min2(end,1), maxtab2 = maxtab2(1:end-1,:); end

% Expand minima boundaries based on threshold

mintab2 = min2(1,:);

for n = 2:length(min2)-1

index = min2(n);

while y2(index) <= min2(n,2)*(1+indicesThreshold), index = index-1;

end

mintemp2(1,:) = [index, y2(index)];

index = min2(n);

while y2(index) <= min2(n, 2)*(1+indicesThreshold)

index = index+1;

if index >= length(y2), index = length(y2); break; end

end

mintemp2(2,:) = [index, y2(index)];

mintab2 = [mintab2 ; mintemp2];

end

mintab2 = [mintab2 ; min2(end,:)];

% Calculate packet metrics

packet_width2 = [];

mean_packet_min2 = [];

packet_integral2 = zeros(1,length(maxtab2));

packet_boundary2 = [];

% Widths and mean minima

for n = 1:2:length(mintab2)-1

packet_width2 = [packet_width2 mintab2(n+1,1)-mintab2(n,1)];

mean_packet_min2 = [mean_packet_min2 mean([mintab2(n+1,2)

mintab2(n,2)])];

end

% Amplitude and integrals

for n = 1:length(maxtab2)

packet_line_x2 = mintab2(n*2-1,1):mintab2(n*2,1);

packet_slope2 = (mintab2(n*2,2)-mintab2(n*2-1,2))/(mintab2(n*2,1)-mintab2(n*2-1,1));

packet_offset2 = mintab2(n*2-1,2) - packet_slope2 * mintab2(n*2-1,1);

packet_line_y2 = packet_line_x2*packet_slope2 + packet_offset2;

packet_boundary2 = [packet_boundary2; [packet_line_x2’

packet_line_y2’]];

packet_amplitude2(n) = maxtab2(n,2) - mean_packet_min2(n);

packet_amplitude_perdiff2(n)=

packet_amplitude2(n)/mean_packet_min2(n);

integral_y2 = y2(mintab2(n*2-1,1):mintab2(n*2,1));

packet_integral2(n) = sum(integral_y2 - packet_line_y2’);

packet_integral_norm2(n) = packet_integral2(n)/mean_packet_min2(n);

end

% Optional figure

if PlotFigs==‘y’

figure;

plot(t,y2,‘linewidth’,2); hold on;

plot(t(maxtab2(:,1)),maxtab2(:,2),‘r.’,…

t(mintab2(:,1)),mintab2(:,2),‘g.’,‘markersize’,15)

plot(t(packet_boundary2(:,1)),packet_boundary2(:,2),‘g.’,‘markersize’,4)

xlabel(‘Seconds’); ylabel(‘Intensity’); title(Name);

end

% Linear fit to minima to estimate drift

packetMin_P = polyfit(t(mintab2(:,1)),mintab2(:,2),1);

min_Slope = packetMin_P(1);

% Summary metrics

packet_frequency2 = length(maxtab2)/(t(mintab2(end,1))-t(mintab2(2,1)))*60;

avg_packet_width2 = mean(packet_width2);

avg_packet_amplitude2 = mean(packet_amplitude2);

avg_packet_amplitude_perdiff2 = mean(packet_amplitude_perdiff2);

avg_packet_integral2 = mean(packet_integral2);

avg_packet_integral_norm2 = mean(packet_integral_norm2);

packet_transport2 = sum(packet_integral2)/(t(mintab2(end,1))-t(mintab2(1,1)))*60;

packet_transport_norm2 = sum(packet_integral_norm2)/(t(mintab2(end,1))-t(mintab2(1,1)))*60;

% Output vector

output = [packet_frequency2 avg_packet_width2 avg_packet_amplitude2 …

avg_packet_integral2 packet_transport2

avg_packet_amplitude_perdiff2 …

avg_packet_integral_norm2 packet_transport_norm2 min_Slope];

end

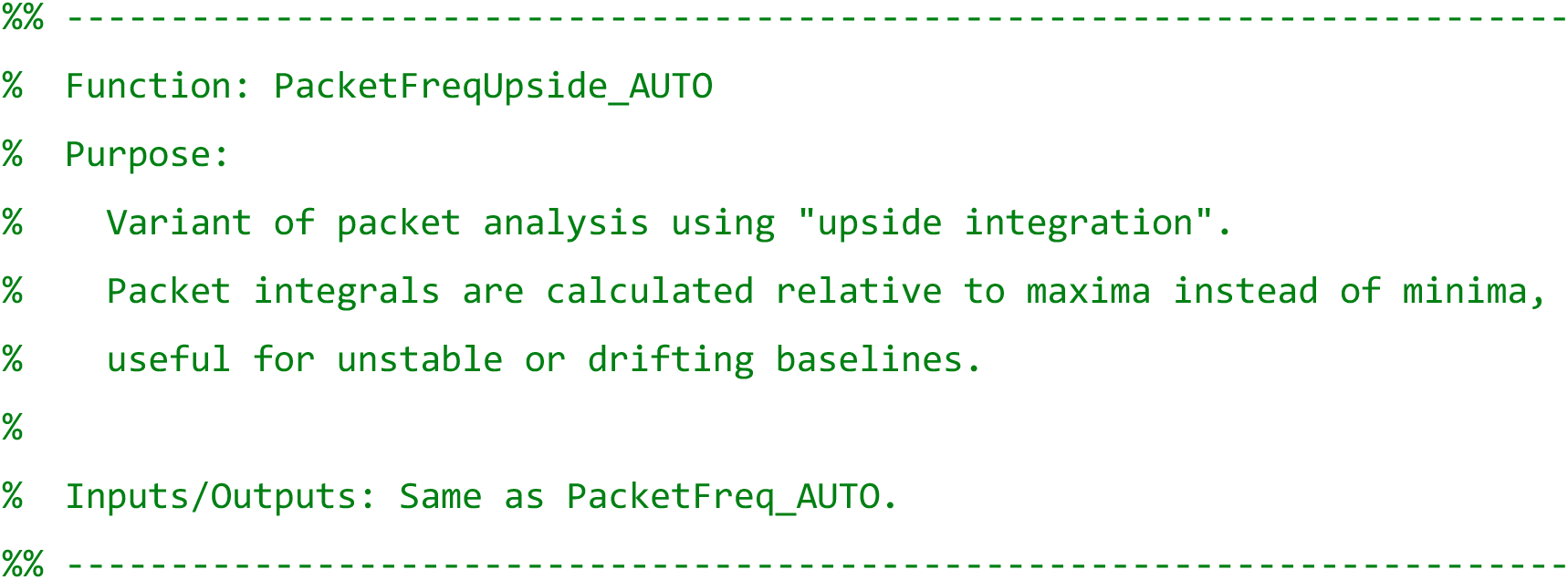

function output = PacketFreqUpside_AUTO(DataFile, thresholds, Name, PlotFigs)

indicesThreshold = thresholds(1);

minThreshold = thresholds(2);

sensitivityThreshold = 1 - thresholds(3);

% Load CSV (time, intensity)

x = csvread(DataFile);

t = x(:,1);

x = x(:,2);

% Smooth trace

y2 = smooth(x,3);

% Adaptive window search

numSoln = [];

minWindow = 1;

maxWindow = length(y2);

for i = minWindow:maxWindow

[x,y] = peakdetYJ(y2, i, minThreshold);

numSoln = [numSoln; length(x)];

end

windowSize = (minWindow:1:maxWindow-1)’;

DiffEQN = movmean(diff(numSoln),10);

OptWindowSize = find(abs(DiffEQN) <= sensitivityThreshold, 1);

[maxtab2, min2] = peakdetYJ(y2, OptWindowSize+1, minThreshold);

if maxtab2(1,1) < min2(1,1), maxtab2 = maxtab2(2:end,:); end

if maxtab2(end,1) > min2(end,1), maxtab2 = maxtab2(1:end-1,:); end

% Expand minima boundaries

mintab2 = min2(1,:);

for n=2:length(min2)-1

index=min2(n);

while y2(index) <= min2(n,2)*(1+indicesThreshold), index=index-1; end

mintemp2(1,:) = [index, y2(index)];

index=min2(n);

while y2(index) <= min2(n,2)*(1+indicesThreshold)

index=index+1;

if index>=length(y2), index = length(y2); break; end

end

mintemp2(2, :) = [index, y2(index)];

mintab2 = [mintab2 ; mintemp2];

end

mintab2 = [mintab2 ; min2(end, :)];

% Packet metrics (relative to maxima)

packet_width2 = [];

mean_packet_min2 = [];

packet_integral2 = zeros(1,length(maxtab2));

packet_boundary2 = [];

for n=1:2:length(mintab2)-1

packet_width2 = [packet_width2 mintab2(n+1,1)-mintab2(n,1)];

mean_packet_min2 = [mean_packet_min2 mean([mintab2(n+1,2)

mintab2(n,2)])];

end

mean_packet_max2 = [];

for n=1:length(maxtab2)-1

mean_packet_max2 = [mean_packet_max2 mean([maxtab2(n,2)

maxtab2(n+1,2)])];

% Linear boundary between minima

packet_line_x2 = mintab2(n*2-1,1):mintab2(n*2,1);

packet_slope2 = (mintab2(n*2,2)-mintab2(n*2-1,2))/(mintab2(n*2,1)-mintab2(n*2-1,1));

packet_offset2 = mintab2(n*2-1,2) - packet_slope2 * mintab2(n*2-1,1);

packet_line_y2 = packet_line_x2*packet_slope2 + packet_offset2;

packet_boundary2 = [packet_boundary2; [packet_line_x2’

packet_line_y2’]];

packet_amplitude2(n) = maxtab2(n,2) - mean_packet_min2(n);

packet_amplitude_perdiff2(n) =

packet_amplitude2(n)/mean_packet_min2(n);

% Upside integration (relative to maxima)

integral_y2 = mean_packet_max2(end) - y2(maxtab2(n,1):maxtab2(n+1,1));

packet_integral2(n) = sum(integral_y2);

packet_integral_norm2(n) = packet_integral2(n)/mean_packet_max2(n);

end

if PlotFigs==‘y’

figure;

plot(t,y2,‘linewidth’,2); hold on;

plot(t(maxtab2(:,1)),maxtab2(:,2),‘r.’,…

t(mintab2(:,1)),mintab2(:,2),‘g.’,‘markersize’,15)

plot(t(packet_boundary2(:,1)),packet_boundary2(:,2),‘g.’,‘markersize’,4)

xlabel(‘Seconds’); ylabel(‘Intensity’); title(Name);

end

% Drift of minima

packetMin_P = polyfit(t(mintab2(:,1)),mintab2(:,2),1);

min_Slope = packetMin_P(1);

% Summary metrics

packet_frequency2 = length(maxtab2)/(t(mintab2(end,1))-

t(mintab2(2,1)))*60;

avg_packet_width2 = mean(packet_width2);

avg_packet_amplitude2 = mean(packet_amplitude2);

avg_packet_amplitude_perdiff2 = mean(packet_amplitude_perdiff2);

avg_packet_integral2 = mean(packet_integral2);

avg_packet_integral_norm2 = mean(packet_integral_norm2);

packet_transport2 = sum(packet_integral2)/(t(maxtab2(end,1))-

t(maxtab2(1,1)))*60;

packet_transport_norm2 = sum(packet_integral_norm2)/(t(maxtab2(end,1))-

t(maxtab2(1,1)))*60;

output = [packet_frequency2 avg_packet_width2 avg_packet_amplitude2 …

avg_packet_integral2 packet_transport2

avg_packet_amplitude_perdiff2 …

avg_packet_integral_norm2 packet_transport_norm2 min_Slope];

end

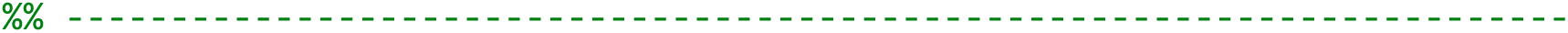
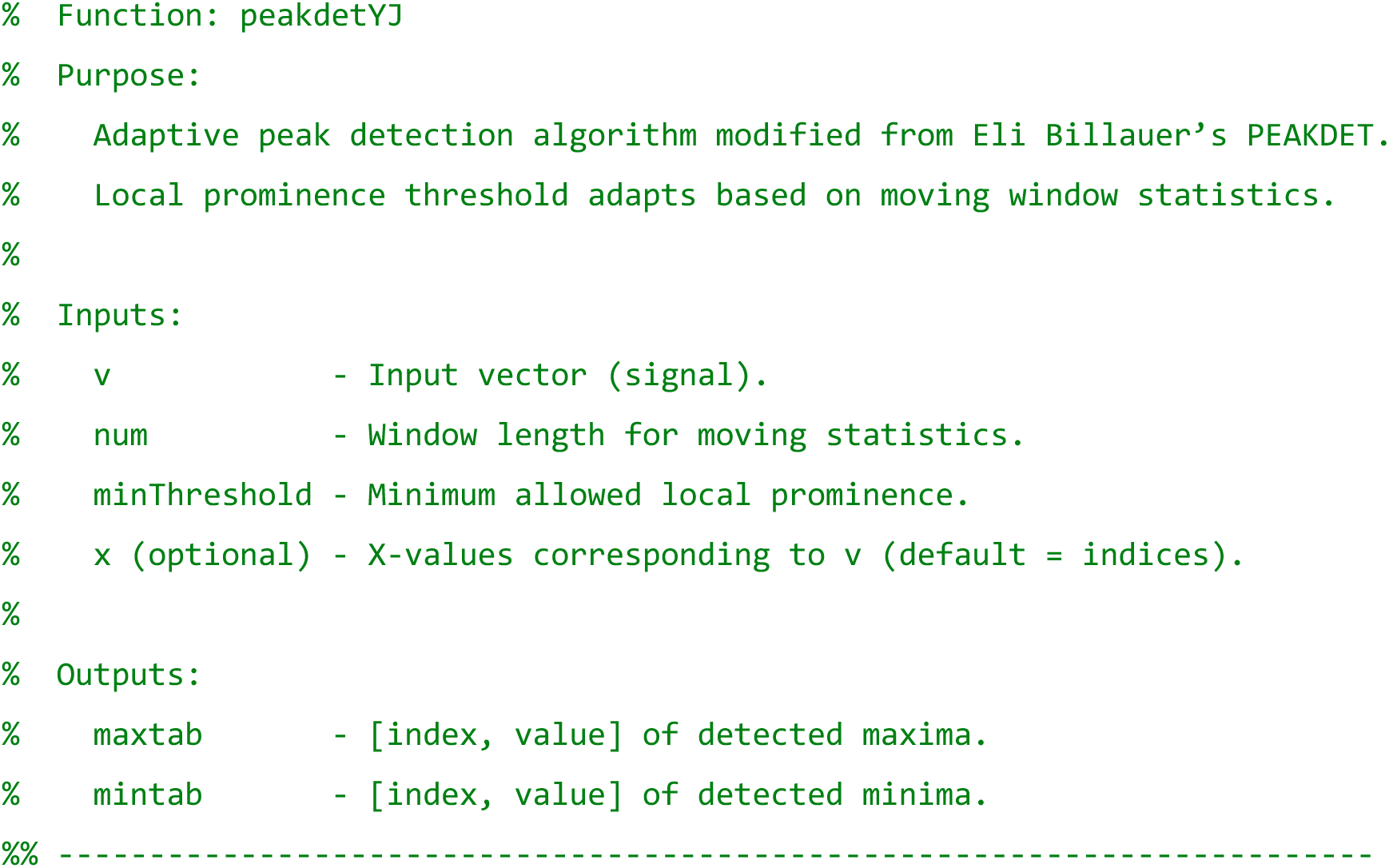

function [maxtab, mintab] = peakdetYJ(v, num, minThreshold, x)

maxtab = [];

mintab = [];

v = v(:);

if nargin < 4

x = (1:length(v))’;

else

x = x(:);

if length(v)∼= length(x)

error(‘Input vectors v and x must have same length’);

end

end

mn = Inf; mx = -Inf;

mnpos = NaN; mxpos = NaN;

lookformax = 1;

% Adaptive local prominence

winLength = num;

winRange = movmax(v,winLength) - movmin(v,winLength);

winRangeMean= movmean(winRange,winLength);

winSTD = movstd(v,winLength);

SNR = 20*log(winRangeMean./winSTD);

LOQ = 10*winSTD./SNR;

prom = 0.2*(winRangeMean-LOQ);

prom(prom<minThreshold) = minThreshold;

critIndex_SNR = find(SNR<15);

prom(critIndex_SNR) = 1.2*prom(critIndex_SNR);

% Iterate through signal

for i=1:length(v)

this = v(i); delta = prom(i);

if this > mx, mx = this; mxpos = x(i); end

if this < mn, mn = this; mnpos = x(i); end

if lookformax

if this < mx-delta

maxtab = [maxtab ; mxpos mx];

mn = this; mnpos = x(i);

lookformax = 0;

end

else

if this > mn+delta

mintab = [mintab ; mnpos mn];

mx = this; mxpos = x(i);

lookformax = 1;

end

end

end

end

